# nanorepertoire: an end-to-end Nextflow pipeline for nanobody repertoire analysis

**DOI:** 10.64898/2026.09.11.750933

**Authors:** Davide Bagordo, Nicola Martelossi, Francesco Lescai

## Abstract

Camelid heavy-chain antibodies, and particularly their variable domains known as nanobodies or VHHs, combine full antigen-binding capacity with a compact and highly stable scaffold, which makes them attractive for both fundamental immunology and therapeutic development. High-throughput adaptive immune receptor repertoire sequencing (AIRR-seq) allows nanobody repertoires to be profiled at great depth, but the analyses applied to VHH data are typically assembled ad hoc from standalone scripts, which limits standardisation and reproducibility across laboratories. Here we present nanorepertoire, an end-to-end Nextflow DSL2 pipeline dedicated to camelid VHH repertoires. It takes paired-end AIRR-seq FASTQ files through quality control, adapter trimming, read merging, in-silico translation, CD-HIT clonotyping and deep-learning CDR3 annotation with nanoCDR-X (Bagordo et al., 2026), and returns an interactive HTML report describing clonal architecture, CDR3 length and amino-acid composition, intra-clonal homogeneity and repertoire diversity, together with the computational carbon footprint of the run. Applied to two publicly available SARS-CoV-2 RBD-selected llama libraries sampled before and after phage-display enrichment (4.8 million paired-end reads in total), the pipeline completed in 59 min on a 16-vCPU cloud instance and recovered 41,363 distinct CDR3 paratopes, reproducing the expected contraction of clonal diversity upon selection. nanorepertoire is open source under the MIT licence at https://github.com/lescailab/nanorepertoire, is archived on Zenodo, and runs unchanged on local, HPC and cloud infrastructures.

## 1. Introduction

The adaptive immune system relies on the generation of diverse receptors to mount highly specific responses against a wide array of antigens. While conventional antibodies comprise both heavy and light chains, camelid species naturally produce heavy-chain antibodies (HCAbs) devoid of light chains. The single variable domain of these antibodies, known as a nanobody or VHH, retains full antigen-binding capacity within a compact, highly stable structure (Muyldermans, 2013). Their distinctive properties, most notably an extended complementarity-determining region 3 (CDR3) capable of accessing cryptic epitopes, render nanobodies extremely promising for therapeutic applications, ranging from viral neutralisation to oncology.

High-throughput adaptive immune receptor repertoire sequencing (AIRR-seq) enables the profiling of nanobody diversity at unprecedented depth, and is now routinely used to mine immune libraries for candidate binders. However, managing the scale and complexity of these data requires robust computational infrastructure. Several established tools exist for conventional antibody and T-cell receptor (TCR) or B-cell receptor (BCR) repertoires, but they rely on germline reference databases and on canonical framework boundaries that transfer poorly to camelid VHH domains. Deep-learning models have recently been developed to resolve the structurally fuzzy boundaries of nanobody CDR loops with high accuracy (Bagordo et al., 2026), yet they currently exist as standalone scripts rather than as components of integrated, end-to-end analytical pipelines. Consequently, laboratories working on VHH repertoires typically assemble custom chains of scripts, which makes these analyses difficult to standardise, to scale and to reproduce across groups. To address these limitations, we developed nanorepertoire, an open-source and scalable Nextflow pipeline. nanorepertoire bridges the gap in nanobodies analyses between raw AIRR-seq data and structural insight, processing sequencing reads into reproducible and interactive repertoire analytics (Fig. 1).

**Fig. 1.**
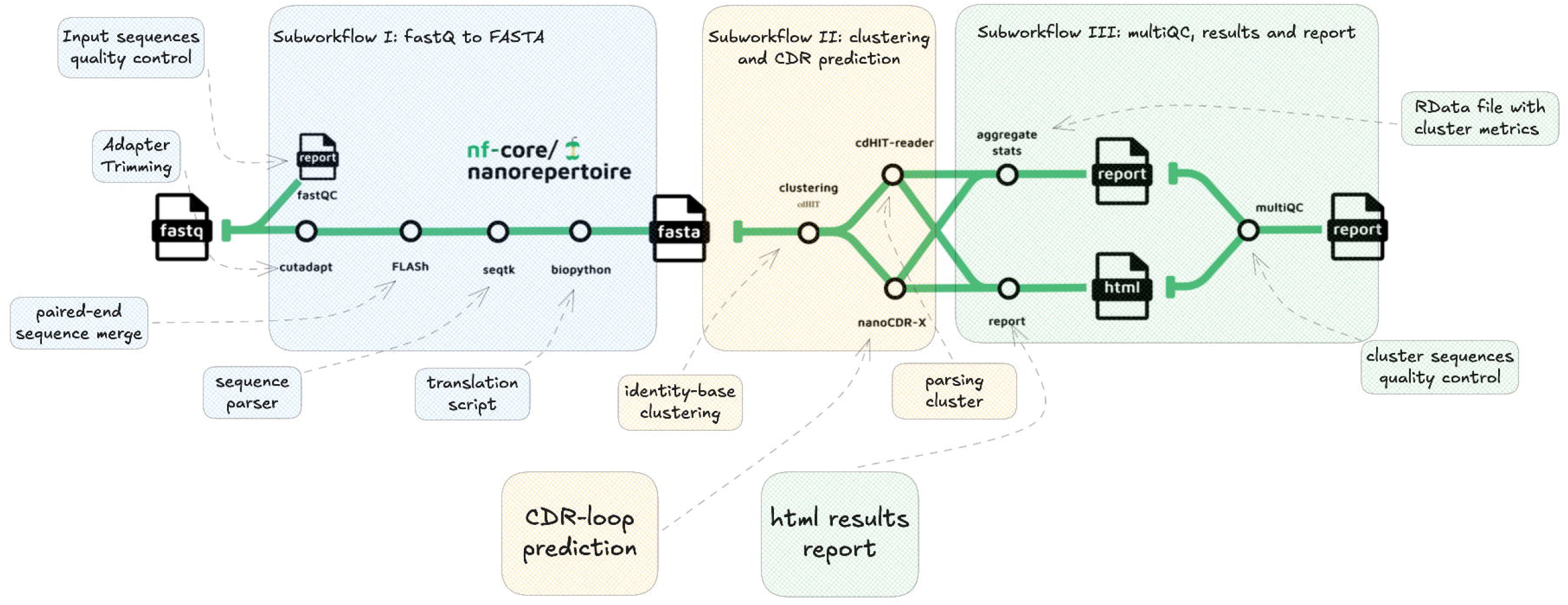
Overview of the nanorepertoire pipeline architecture. The workflow is divided into three sequential subworkflows: sequence pre-processing (Subworkflow I), clustering and CDR prediction (Subworkflow II), and quality control and reporting (Subworkflow III). The CDR-loop prediction module (nanoCDR-X) and the HTML results report constitute the core analytical engine of the pipeline. *Alt text*. Flow diagram of the three nanorepertoire subworkflows, from raw FASTQ files to the HTML report.

## 2. Materials and methods

### 2.1. Implementation and execution

nanorepertoire is implemented in Nextflow DSL2 (Di Tommaso et al., 2017). Portability across computational environments, local workstations, HPC clusters and cloud infrastructures is handled by Nextflow executors and configuration profiles, so that the same workflow definition runs unchanged on all of them. Software dependencies are resolved through containers, and the repository ships ready-to-use docker and singularity profiles; a Conda profile is also provided as a fallback for environments in which containers cannot be used. The workflow has been validated with Nextflow 25.04.0. The complete list of tools and versions used by the pipeline is given in Supplementary Table S3. The workflow is modularised into three subworkflows (Fig. 1).

### 2.2. Pre-processing (fastq_to_fasta)

The pipeline accepts raw paired-end FASTQ files. Initial quality control is performed with FastQC, followed by adapter trimming with Cutadapt (Martin, 2011). Paired-end reads are merged with FLASH (Magoč and Salzberg, 2011) to reconstruct the full-length VHH sequences, and sequence formatting and standardisation are handled with seqtk and Biopython. Sequences are subsequently translated from nucleotide to amino acid with an integrated custom tool (Nanotranslate).

### 2.3. Clustering and annotation (fasta_clustering)

To define clonotypes, the translated amino-acid sequences are clustered at a configurable sequence-identity threshold (90% by default) with CD-HIT (Li and Godzik, 2006). This step reduces data redundancy and identifies clonally expanded populations. The resulting clusters are then processed to extract structural annotations: the pipeline integrates nanoCDR-X (Bagordo et al., 2026), a deep-learning framework dedicated to the precise identification and extraction of CDR3 regions from full-length nanobody sequences. This step is essential to resolve the structural fuzziness of VHH binding loops and to enable accurate downstream functional inference.

### 2.4. Reporting and metrics (repertoire_report)

A defining feature of nanorepertoire is its dedicated reporting architecture. Quality metrics from the pre-processing steps are aggregated with MultiQC (Ewels et al., 2016). The biological data, including cluster sizes, CDR3 lengths and sequence distributions, are processed through an R-based statistical engine (aggregate_stats) that outputs serialised objects and tabular summaries. A Python/Plotly module then generates a comprehensive, interactive HTML dashboard offering dynamic visualisations of clonal expansion and repertoire diversity (Fig. 2). In addition, the pipeline natively incorporates the nf-co2footprint plugin to monitor and report the computational carbon emissions and energy usage of each execution, following the Green Algorithms method (Lannelongue et al., 2021) and promoting sustainable bioinformatics practices.

**Fig. 2.**
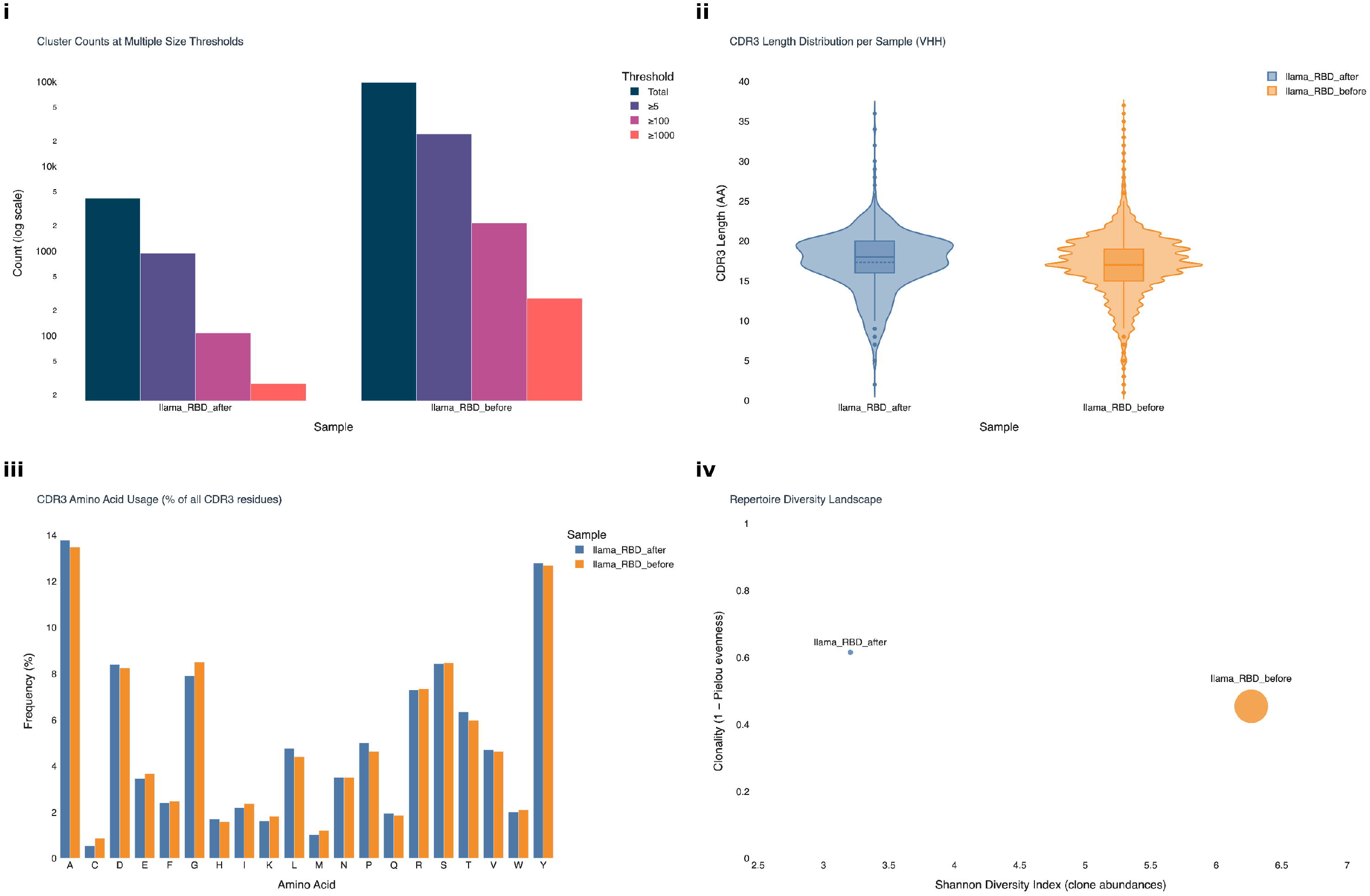
Representative panels of the interactive HTML report generated by nanorepertoire (complete report in Supplementary File S1). (i) Cluster counts at four cumulative size thresholds (total, ≥5, ≥100, ≥1000 members) on a logarithmic axis. (ii) Per-sample CDR3 length distribution as a violin with embedded box plot. (iii) CDR3 amino-acid usage as a percentage of all CDR3 residues. (iv) Repertoire diversity landscape: Shannon index of clone abundances against clonality (1 ™ Pielou evenness), with marker area proportional to the number of unique CDR3 paratopes. All panels are produced automatically by the workflow. *Alt text*. Four charts from the nanorepertoire report: a logarithmic bar chart of cluster counts at four size thresholds, a violin plot of CDR3 lengths per sample, a grouped bar chart of amino-acid usage across the twenty residues, and a bubble chart of Shannon diversity against clonality.

## 3. Results

### 3.1. Benchmark dataset and computational performance

To demonstrate the workflow end to end, we processed a publicly available pair of SARS-CoV-2 RBD-selected llama VHH libraries (Xu et al., 2021; SRA accessions SRR13768393 and SRR13768392), sampled before and after phage-display enrichment, for a combined total of 4,771,793 raw paired-end reads. The two libraries were not sequenced to comparable depth: 4,376,579 read pairs before enrichment against 395,214 after. This asymmetry is itself instructive, in that it exercises the depth-normalised statistics the report derives for this specific scenario. The analysis was run on a single dedicated Google Cloud instance (n2-standard-16; 16 vCPUs, 64 GB RAM) with the Nextflow local executor, which dispatched the 19 processes of the workflow as independent tasks scheduled concurrently on that machine according to their declared resource requirements. It completed from raw FASTQ to rendered report in 58 min 42 s of wall-clock time and 6.5 cumulative CPU-hours, the rate-limiting steps being deep-learning CDR3 annotation (24 min) and CD-HIT clustering (17 min) of the larger library; the integrated nf-co2footprint plugin estimated the environmental cost at 67 g CO_2_e (141 Wh). Across the two libraries, 94,431 translated sequences received a CDR3 annotation, 102,593 clonotypes were resolved at the 90% identity threshold and 41,363 distinct CDR3 paratopes were recovered, with a mean length of 17.1 amino acids (SD 3.8; median 17). The complete report is provided as Supplementary File S1.

### 3.2. Clonal cluster analysis

The first section of the report presents a per-sample metrics table followed by three views of clonal architecture. The table exposes clonotype richness, clustered sequences, depth-normalised richness, Shannon and Simpson indices on clone abundances, clonality, the Gini coefficient, D50, and the fraction of the library held by the largest and by the ten largest clonotypes: here 98,399 clonotypes over 3,210,691 clustered sequences before enrichment against 4,194 over 324,341 after, or 30.7 and 12.9 clonotypes per thousand sequences once normalised for depth.

The clonal expansion profile stratifies clonotypes into four size classes (fewer than 5, 5 to 99, 100 to 999, and 1000 or more members) and recovers in both libraries the skewed architecture characteristic of an immune repertoire, with 75.5% and 77.6% of clonotypes in the smallest class. The cluster-abundance panel (Fig. 2i) presents the same stratification cumulatively on a logarithmic axis, so that four orders of magnitude are legible at once: the decay from 98,399 to 24,093, 2,127 and 277 clonotypes before enrichment, against 4,194 to 938, 108 and 27 after, makes the steepness of each clonal hierarchy comparable between samples of very different size. The top ten panel resolves the extreme tail, listing each dominant clonotype with the identifier of its representative sequence: here the largest post-enrichment clonotype comprised 120,010 sequences, 24-fold the tenth-ranked one, and the ten largest accounted for 75.2% of that library against 40.2% before. Each entry is keyed to a representative identifier in clusterbig.csv, from which a screening campaign can proceed directly.

### 3.3. CDR3 diversity analysis

The second section characterises the annotated paratopes. A per-sample count of distinct CDR3 sequences recorded 40,089 unique paratopes before enrichment and 1,274 after; like every raw count, these scale with sequencing depth, and the report therefore presents them alongside the depth-normalised statistics of the preceding section rather than as a diversity measure.

The length distribution is shown as a violin with an embedded box plot (Fig. 2ii), which reports the median, interquartile range and full spread per sample in a single mark: median 17 residues (IQR 15–19, range 1–37) before enrichment and 18 residues (IQR 16–20, range 2–36) after. The frequency profile renders the same distributions as overlaid curves over lengths 0–45, a view better suited to detecting multiple modes and to comparing the shoulders of two samples directly; the modal length was 17 residues before and 20 after, with 59.3% and 64.5% of paratopes in the 17–23 residue window and 8.5% below 12 residues in both libraries.

CDR3 boundaries are called by nanoCDR-X, the deep-learning annotator we developed for camelid VHH domains, and every run benchmarks it against the regular-expression definition used previously for nanobody repertoires (Deschaght et al., 2017). Where both apply, agreement is essentially complete: 99.3% and 99.2% of sequences match exactly, with a median offset of zero at both termini.

### 3.4. CDR3 amino-acid composition

The third section profiles the chemistry of the annotated paratopes with two complementary views of the same matrix. A heat map gives residue frequency as a percentage of all CDR3 positions, with samples as rows and residues as columns, and is the more compact view as the number of libraries grows; a grouped bar chart (Fig. 2iii) places the samples side by side per residue, which is the view that permits a difference at a single position to be read off directly.

In this run, six residues accounted for approximately 58.7% of all CDR3 positions in both libraries: alanine (13.5–13.8%), tyrosine (12.7–12.8%), serine (8.4–8.5%), aspartate (8.3–8.4%), glycine (7.9–8.5%) and arginine (7.3%), reproducing the aromatic- and hydroxyl-rich paratope chemistry documented for camelid VHH domains and thereby providing an internal consistency check on the annotation step. The only residue to differ appreciably between conditions was cysteine, at 0.86% before enrichment and 0.54% after.

### 3.5. Clonality and intra-clonal homogeneity

The final section summarises variation within clonotypes. For each cluster, the report takes the mean identity of its non-representative members to the representative and plots that quantity across clusters; singletons are excluded, being undefined. Among the 49,723 and 2,217 clusters for which it is defined, 90.1% and 82.4% fell below full identity, with median identities of 95.3% and 97.4%. The distribution is bounded below by the clustering threshold, and the panel states this truncation explicitly: it describes homogeneity only within that window and is measured against an empirically chosen cluster member rather than an inferred germline, so it is descriptive rather than a germline-referenced mutation rate.

The diversity landscape (Fig. 2iv) closes the report, placing each library on abundance-based axes, the Shannon index of clone abundances against clonality, with marker area proportional to the number of unique paratopes. The two libraries separated on both axes, from H’= 6.27 and clonality 0.455 before enrichment to H’= 3.21 and clonality 0.616 after.

## 4. Conclusion

nanorepertoire provides the nanobody community with a portable and reproducible route from raw AIRR-seq reads to interpretable repertoire analytics. It integrates deep-learning CDR3 annotation into a standardised workflow and reports, in a single automated document, the quality of that annotation, the diversity statistics that depend on it, and the environmental cost of obtaining them.

## Supporting information

Example Report

## CRediT authorship contribution statement

**Davide Bagordo:** Conceptualization, Methodology, Software, Formal analysis, Investigation, Visualization, Writing – original draft. **Nicola Martelossi:** Validation, Writing – review & editing. **Francesco Lescai:** Conceptualization, Supervision, Funding acquisition, Resources, Writing – review & editing.

## Data availability

The sequencing data analysed in this study are publicly available from the NCBI Sequence Read Archive under accessions SRR13768392 and SRR13768393. The source code of nanorepertoire is openly available at https://github.com/lescailab/nanorepertoire under the MIT licence, and the version used in this work is archived on Zenodo (https://doi.org/10.5281/zenodo.17379841).

