## Supplementary material for "nanorepertoire: an end-to-end Nextflow pipeline for nanobody repertoire analysis": Example Report

### Nanorepertoire

Nanobody Repertoire Analysis (VHH Sequencing · Clonotype & CDR3 Analysis Report)

**Pipeline Execution Metrics**

If you are interested in detailed Start/End Times, CPU Efficiency, and $CO_2$ metrics, please see the dedicated reports produced upon completion of the pipeline. We sincerely thank the authors of the

**nf-co2footprint plugin**

and the Green Algorithms project for enabling these measurements.

**Report Generated**

July 27, 2026

#### Summary

SEQUENCES WITH A CALLED CDR3

**94,431**

Total across all samples

TOTAL CLUSTERS (CD-HIT)

**102,593**

90% identity threshold

UNIQUE CDR3 SEQUENCES

**41,363**

Distinct paratopes

MEAN CDR3 LENGTH

**17.1 AA**

SD: 3.8 · Median: 17

SAMPLES ANALYSED

**2**

Independent libraries

#### 1 Clonal Cluster Analysis

Amino acid sequences were clustered using **CD-HIT** at 90% identity, following the landmark approach of Deschaght et al. (2017). **Expanded clonotypes** (≥5 members) represent B-cell lineages that have undergone antigen-driven somatic expansion. Large clusters (≥1000 members) are dominant clonal responses — prime candidates for high-affinity nanobodies. The cluster size distribution mirrors the "clonotype expansion" readout of tools like MiXCR and IMGT/VQuest.

*Deschaght et al. 2017. Front. Immunol. doi:10.3389/fimmu.2017.00420 · Bolotin et al. 2015 (MiXCR). Nature Methods.*

| Sample | Unique CDR3s | Clusters | Clustered sequences | Clusters / 1k seqs | Shannon H' (abundance) | Clonality | S |
|---|---|---|---|---|---|---|---|
| **llama_RBD_after** | 1,274 | 4,194 | 324,341 | 12.93 | 3.205 | 0.616 | [illegible] |
| **llama_RBD_before** | 40,089 | 98,399 | 3,210,691 | 30.65 | 6.268 | 0.455 | [illegible] |

#### Clonal Expansion Profile

Breakdown of clusters by size class per sample. Singletons represent rare or noisy sequences. Expanded clones (≥5 members, orange/red) indicate B-cell lineages selected by antigen exposure.

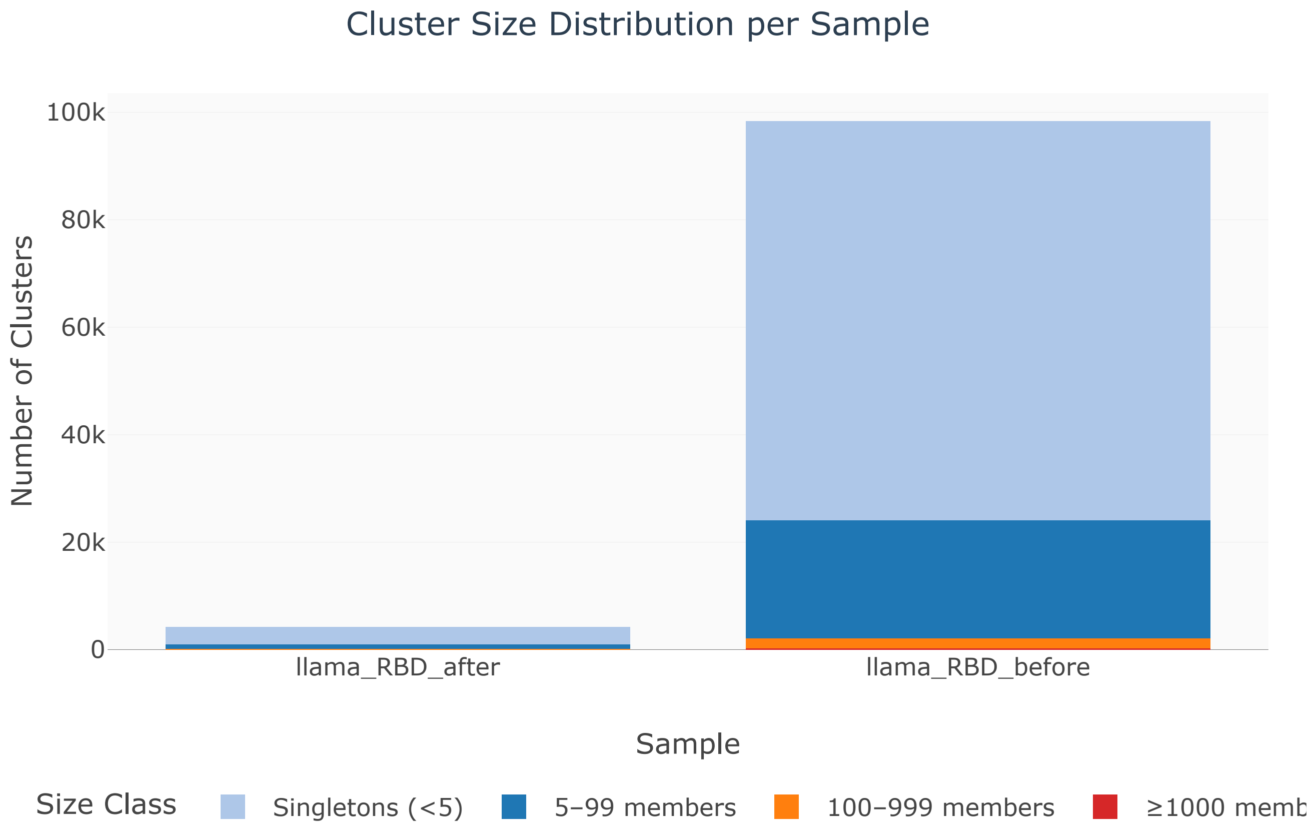

#### Cluster Abundance at Size Thresholds

Cluster counts at four size thresholds (log scale). Reveals the hierarchical repertoire structure from singletons to dominant clones.

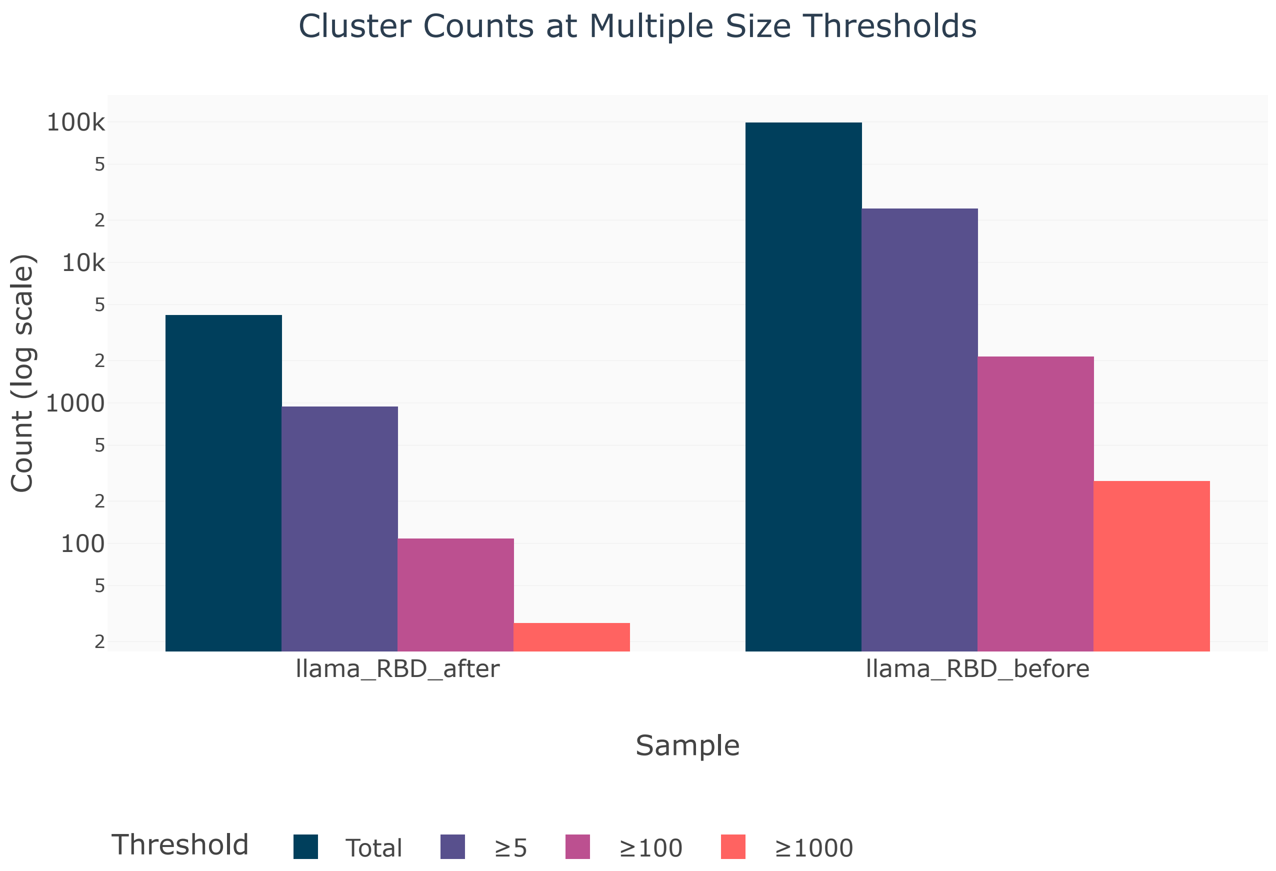

##### Top-10 Largest Clonotypes per Sample

The 10 most abundant clusters per sample. Dominant clones in immunized animals likely represent the most affinity-matured antigen binders.

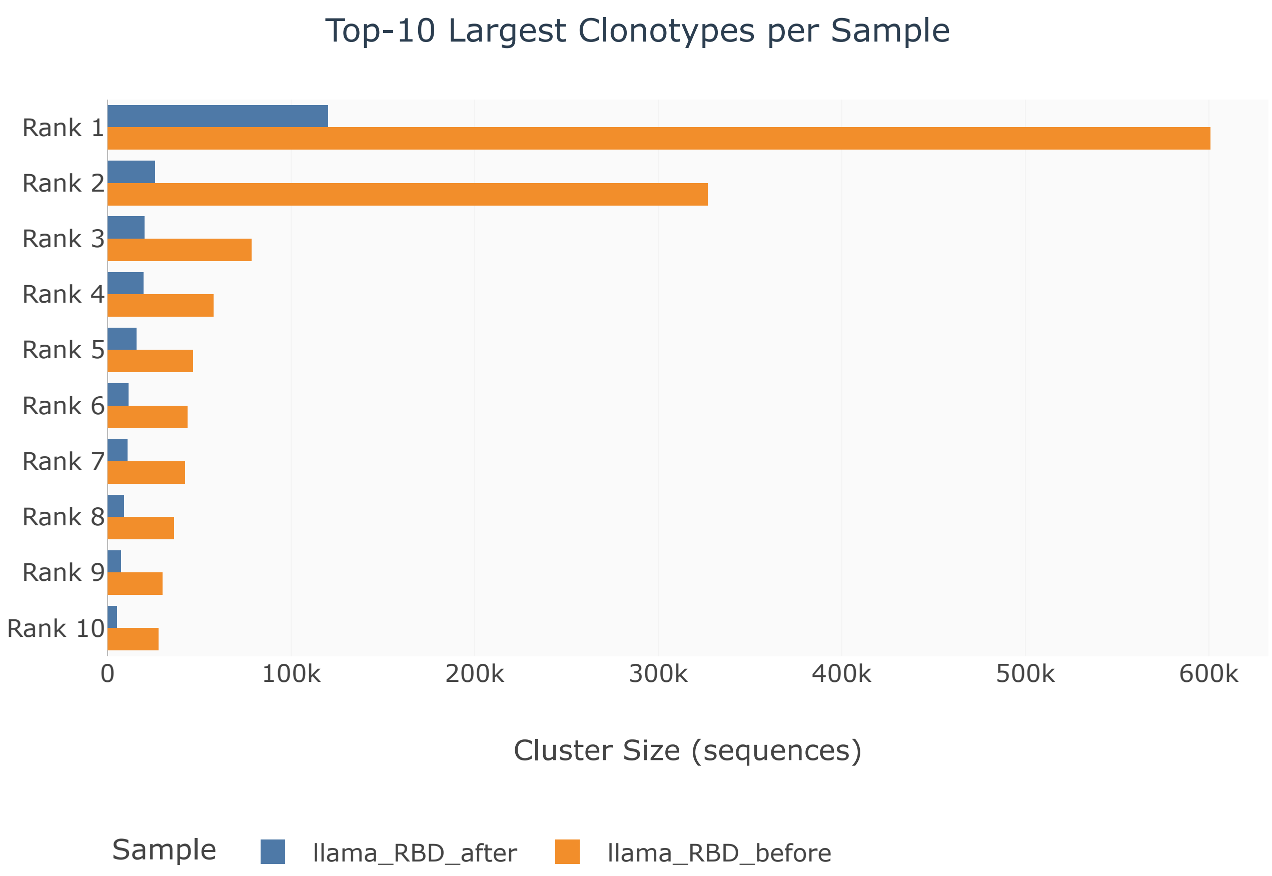


#### 2 CDR3 Diversity Analysis

The **CDR3** (Complementarity-Determining Region 3) is the primary antigen-binding loop of VHH nanobodies. VHH CDR3s are typically *longer and more diverse* than VH CDR3s in conventional antibodies (VHH mean ~19 AA vs VH ~12 AA — Spinelli et al. 2022), enabling access to deep epitope cavities and enzyme active sites. CDR3 length diversity correlates with repertoire breadth; antigen-enriched samples may show length skewing toward specific structural motifs. Shorter CDR3s (<12 AA) tend to form flat β-strand paratopes; longer CDR3s (>16 AA) adopt protruding finger-like conformations.

*Spinelli et al. 2022. Front. Immunol. doi:10.3389/fimmu.2022.927966 · Muyldermans 2013. Annu. Rev. Biochem.*

##### Unique CDR3 Sequences per Sample

Total number of distinct CDR3 sequences per sample. Higher numbers indicate broader repertoire diversity.

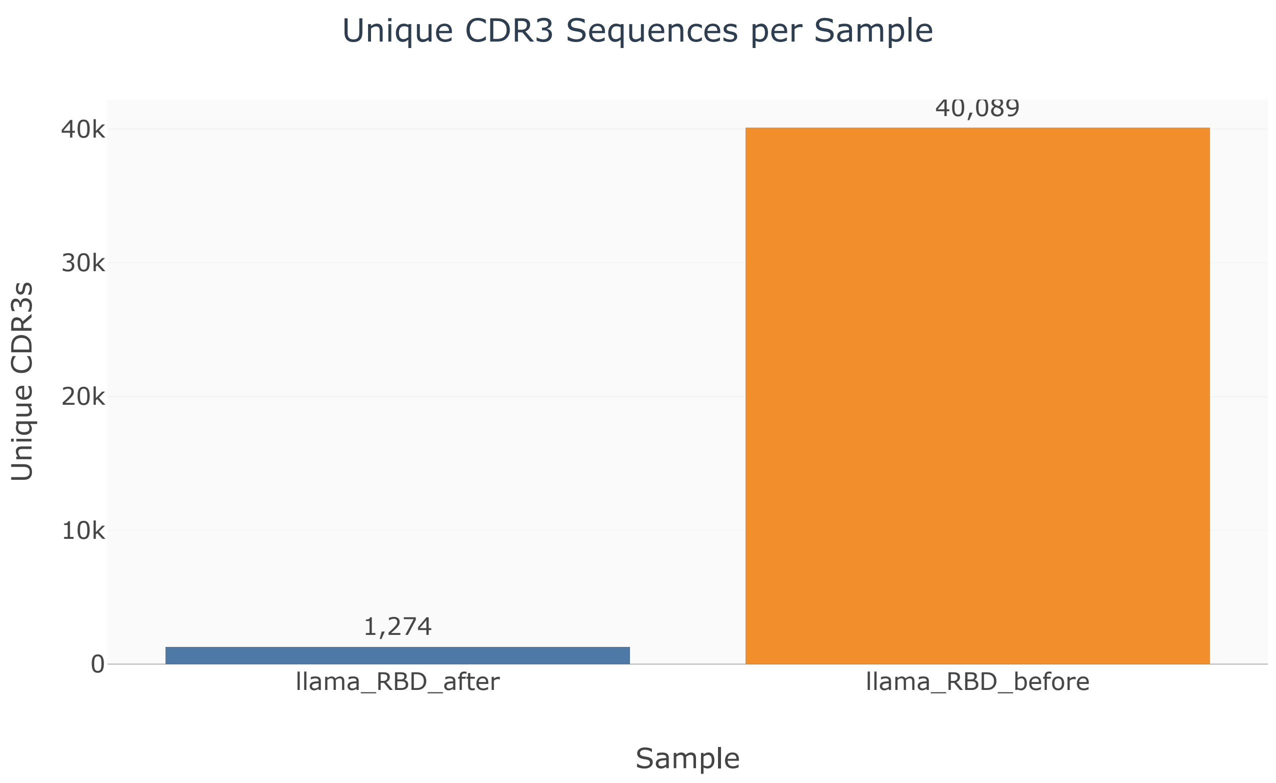

##### CDR3 Length Distribution (Violin)

Violin plot of CDR3 length distribution per sample. Box shows IQR; line shows median. VHH CDR3s typically peak at 15–23 AA.

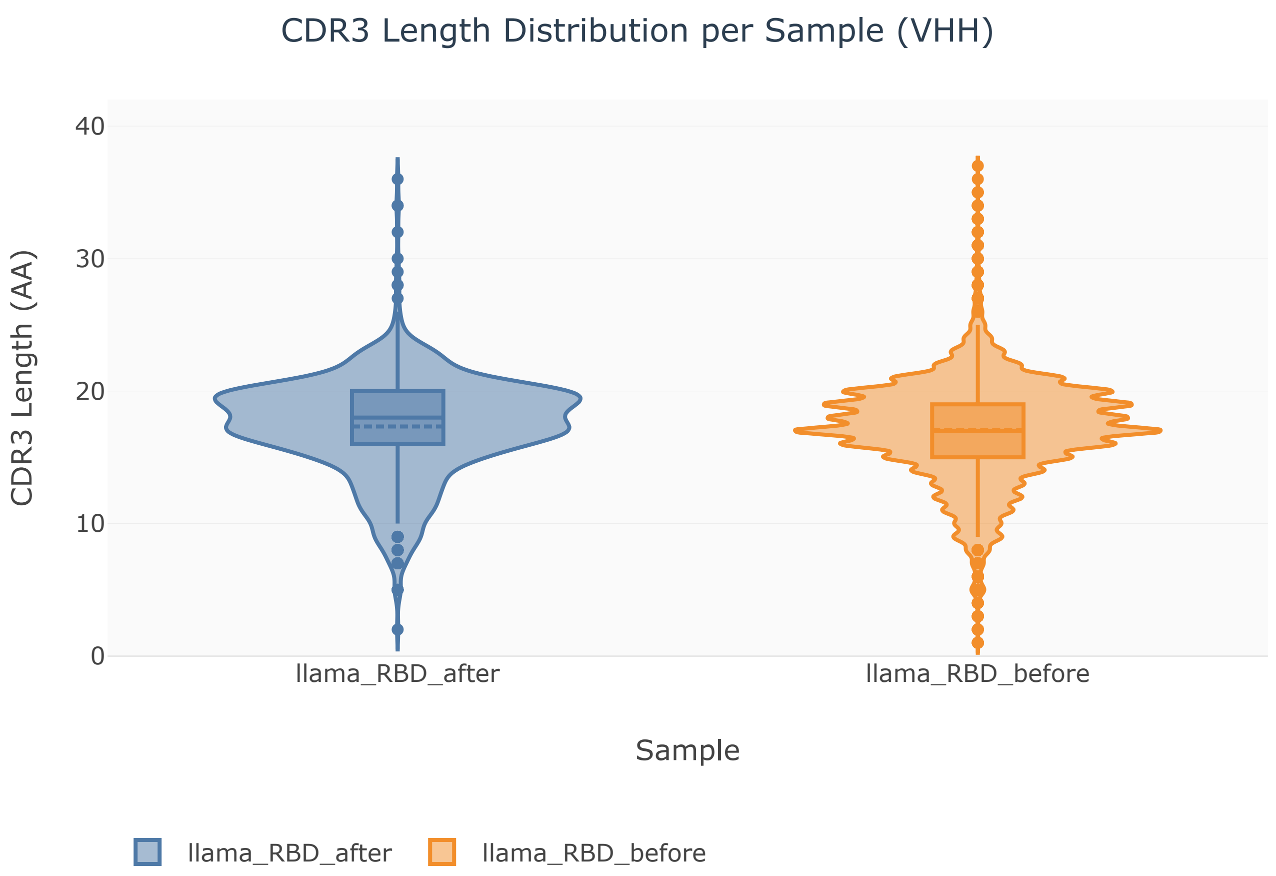

##### CDR3 Length Frequency Profile

Area chart of CDR3 length frequency per sample. Peaks at specific lengths may indicate dominant structural motifs or antigen-driven convergence. Short CDR3s (<12 AA): flat β-strand paratopes. Long CDR3s (>16 AA): protruding loops for cavity/groove binding.

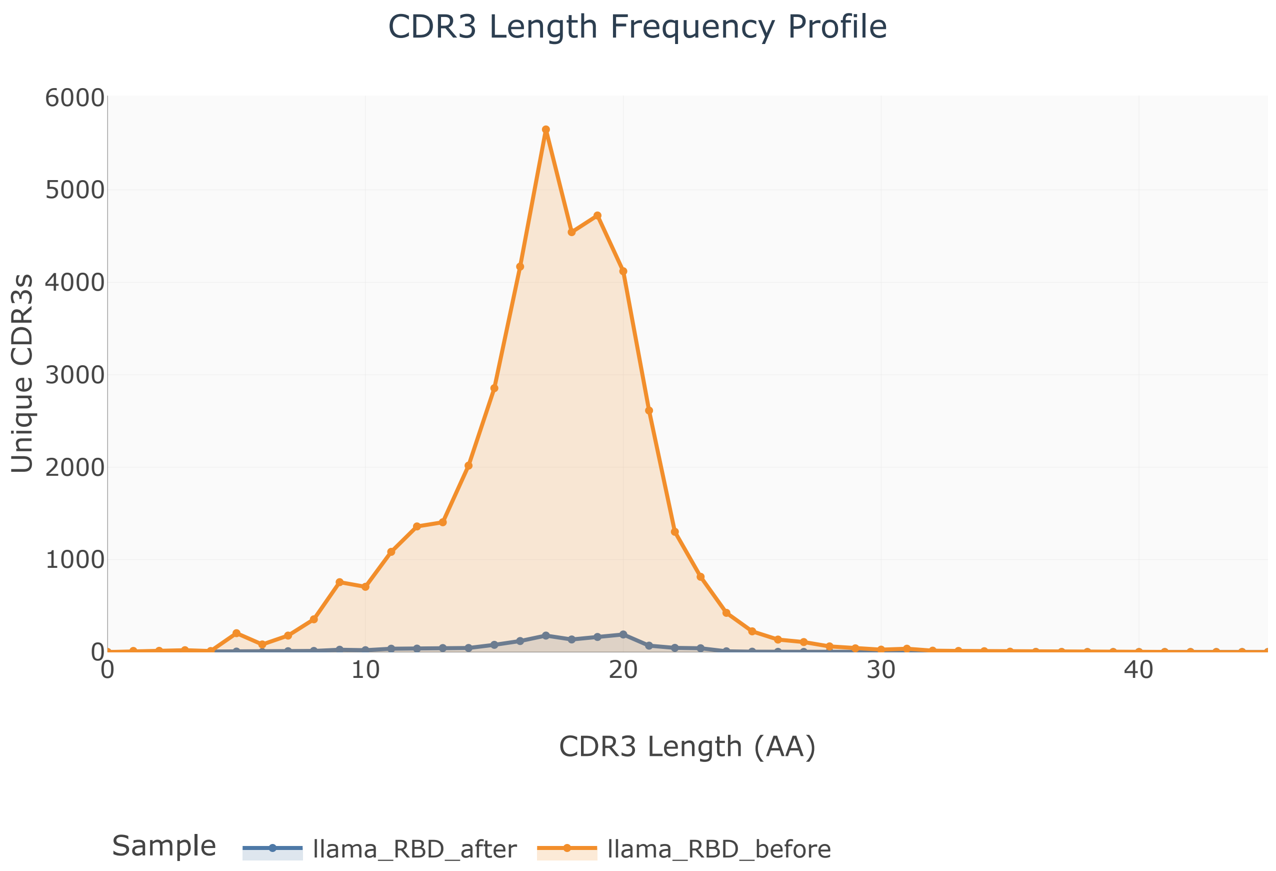


#### CDR3 annotation QC

> Every length and composition figure above depends on where the CDR3 boundaries were placed, so the calls of the deep-learning annotator are compared here against the motif-based definition documented in the Methods: the loop starts immediately after the cysteine of the framework-3 anchor (`T..Y.C`, the `YYC` motif) and ends immediately before the tryptophan of the framework-4 anchor (`WG.G`, the `WGQ` motif; reads in which that tryptophan is substituted are anchored on the C-terminal `TVSS` motif instead). An offset of 0 at both ends means the two definitions agree residue for residue. A systematically *positive* C-terminal offset would mean the annotated CDR3 extends into framework 4, which is rich in A, T and V, and would inflate those residues in the composition panels. Sequences in which neither anchor can be located are not comparable and are counted separately. The full offset distribution is written to `cdr3_boundary_qc.tsv` and `cdr3_boundary_offsets.tsv` next to this report.

| Sample | Sequences compared | Exact agreement | Median N-terminal offset | Median C-terminal offset | Median called length | Median reference length |
|---|---|---|---|---|---|---|
| **llama_RBD_after** | 1,563 / 1,687 | 99.23% | 0.0 | 0.0 | 18.0 | 18.0 |
| **llama_RBD_before** | 86,372 / 92,752 | 99.31% | 0.0 | 0.0 | 17.0 | 17.0 |

##### CDR3 Boundary Offsets

Distribution of the offsets between the called and the motif-based boundaries, per sample and per terminus, as a percentage of the sequences that could be compared. The y axis is logarithmic so that rare disagreements remain visible next to the dominant zero-offset bar.

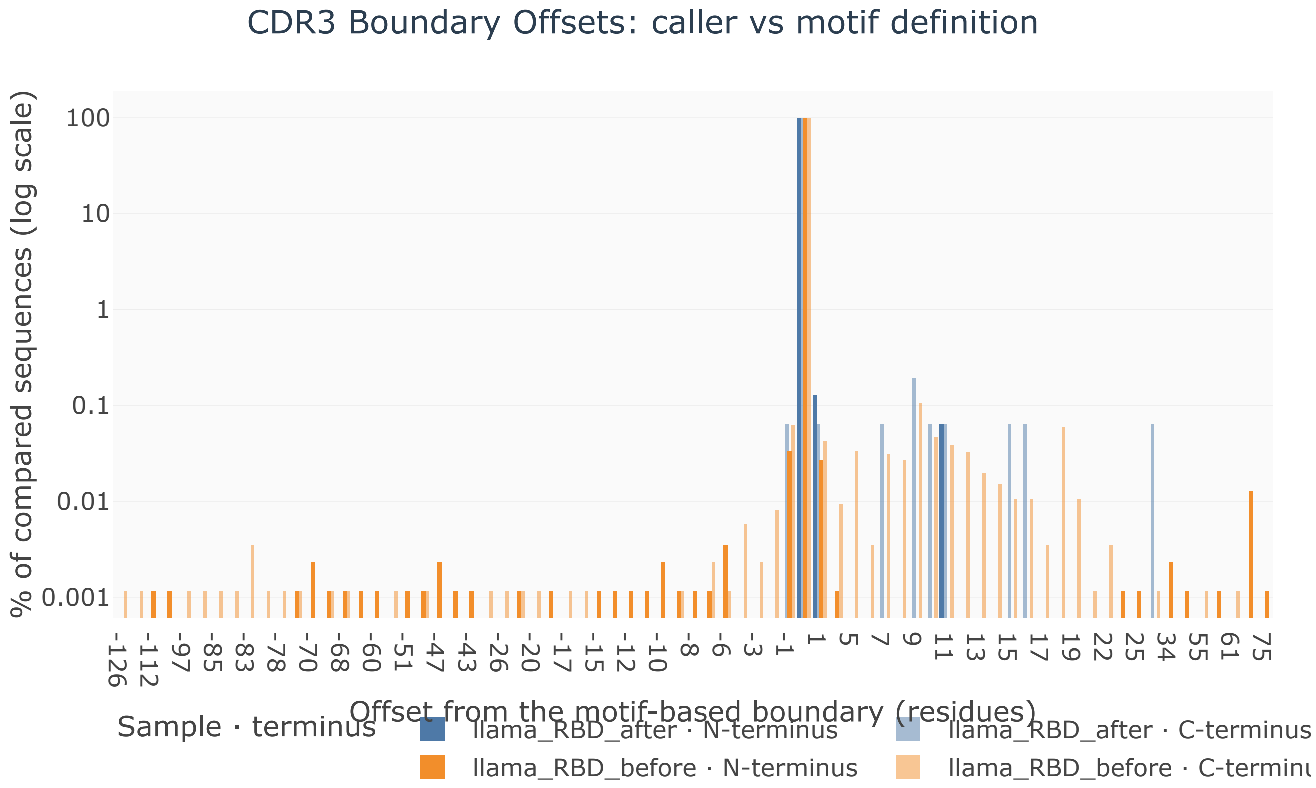

### 3 CDR3 Amino Acid Composition

The amino acid (AA) composition of CDR3 loops reflects antigen-driven selection. VHH CDR3s are characteristically enriched in **Tyrosine (Y)** and **Serine (S)** — enabling hydrogen bonding and Van der Waals contacts. **Cysteine (C)** pairs can form intra-CDR3 disulfide bonds with CDR1, creating a structural scaffold that extends paratope reach. Charged residues (R, D, E, K) mediate electrostatic complementarity. Convergent AA enrichment across samples suggests shared antigen pressure.

*Desmyter et al. 2002. J. Biol. Chem. doi:10.1074/jbc.D200025200 · Mitchell & Colwell 2018. Proteins.*

##### Amino Acid Frequency Heatmap

Heatmap of amino acid frequency (% of all CDR3 residues) per sample. Dark = high frequency. Key VHH residues: Y, S, G, T. Cysteine (C) involvement suggests disulfide-mediated loop stabilization.

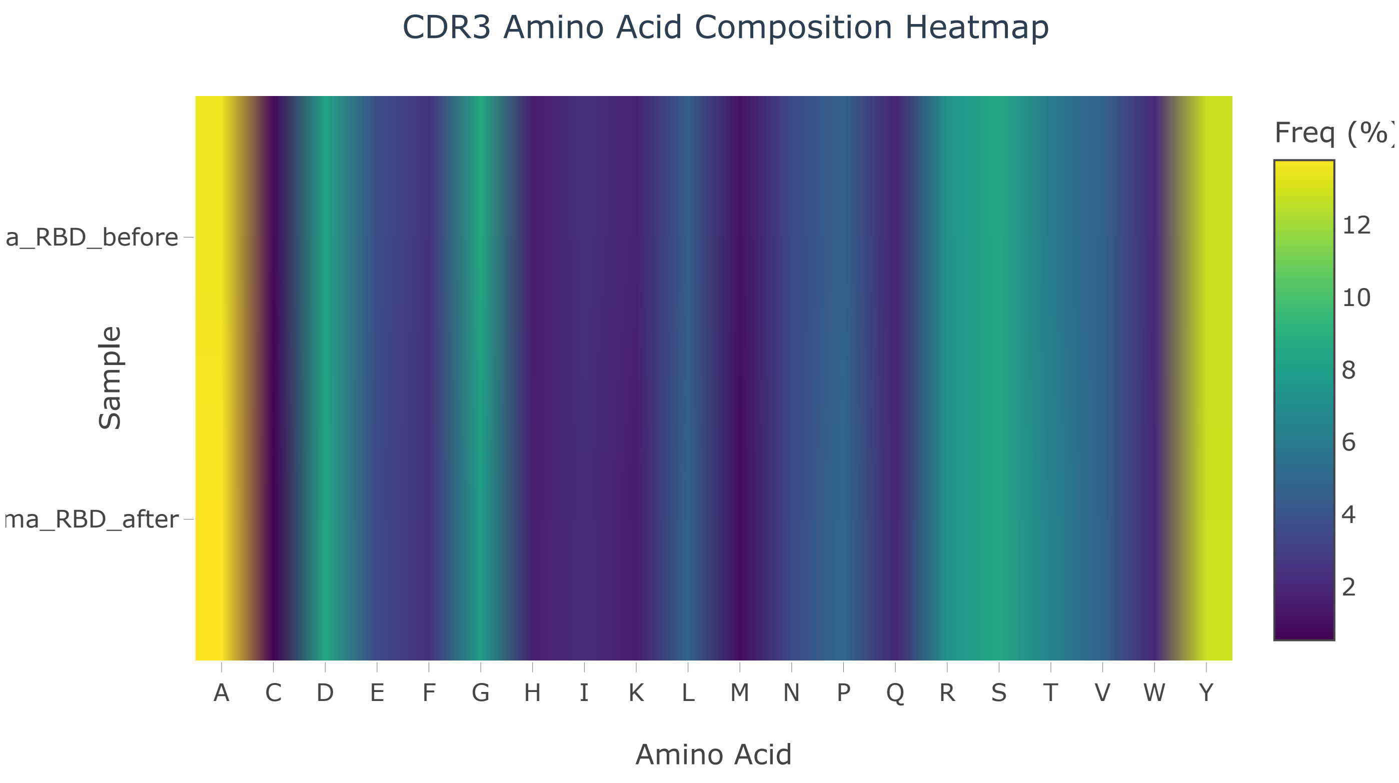

##### Side-by-Side Amino Acid Usage

Side-by-side comparison of amino acid usage per sample. Differences across samples may reflect divergent antigen-binding strategies or B-cell selection biases.

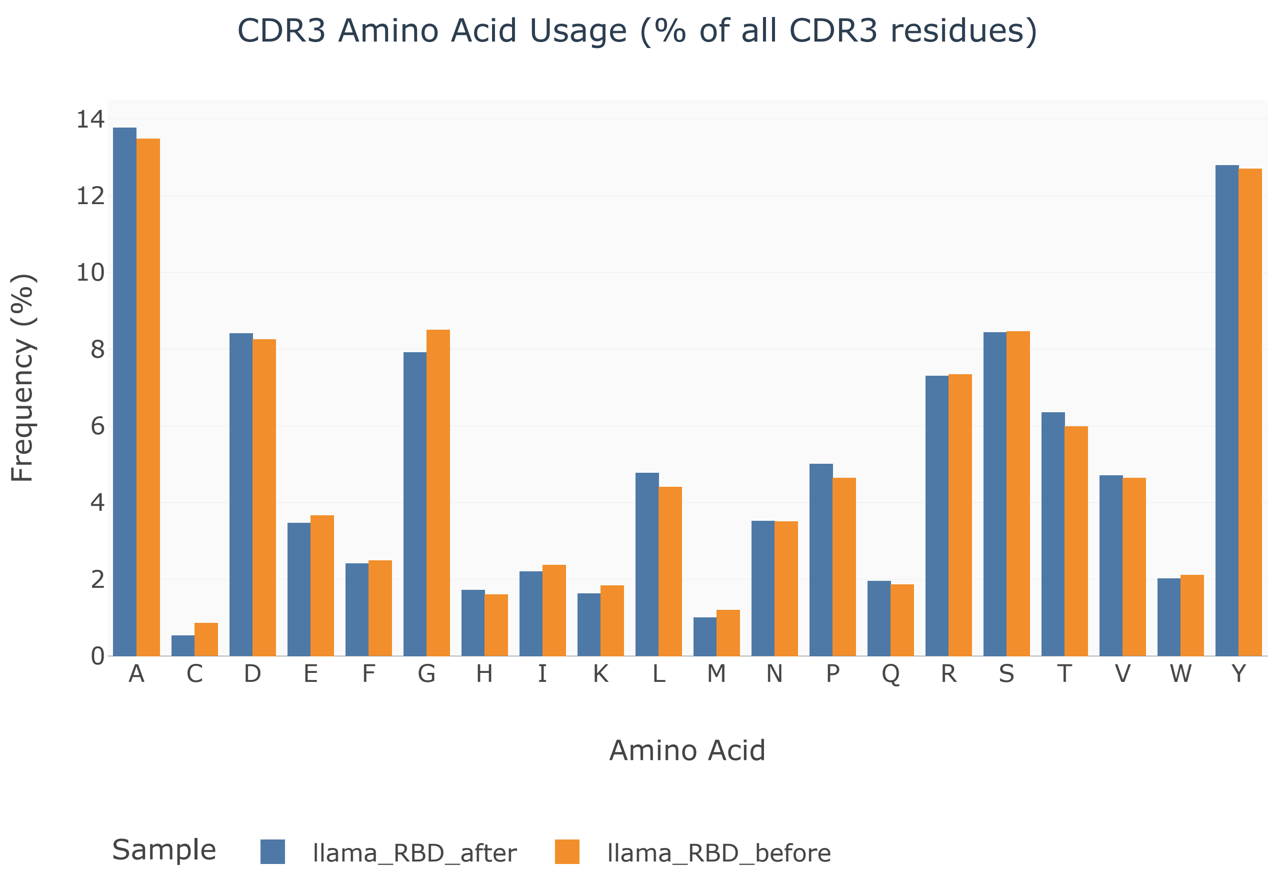

#### 4 Clonality & Intra-clonal Homogeneity

The mean identity of the members of a cluster to its representative is a **descriptive measure of intra-clonal homogeneity**: clusters close to 100% are internally uniform, clusters closer to the clustering threshold group sequences that differ more from their representative. **This quantity is truncated by construction**: CD-HIT admits a sequence into a cluster only if its identity to the representative is at least 90%, so no value below that threshold can be observed and the distribution is bounded on the left by the clustering parameter itself. It is therefore *not* an estimate of somatic hypermutation, and it is not equivalent to the V-gene mutation frequency reported by MiXCR or IMGT/V-QUEST, which is measured against an inferred germline reference rather than against an empirically chosen cluster member; the analogy is conceptual only. Sequence-level differences within a cluster may equally reflect somatic mutation, PCR and sequencing error, or repeated sampling of the same molecule, which this workflow does not deduplicate and therefore cannot tell apart.

*Li & Godzik 2006 (CD-HIT). Bioinformatics. doi:10.1093/bioinformatics/btl158 · Bolotin et al. 2015 (MiXCR). Nature Methods. doi:10.1038/nmeth.3364*

##### Mean Member Identity to Cluster Representative

Distribution of the mean identity of cluster members to their representative, one value per cluster (n = 51,940 multi-member clusters; singletons are excluded because the quantity is undefined for them). Values are bounded below by the 90% clustering threshold, so the panel describes homogeneity within that window only.

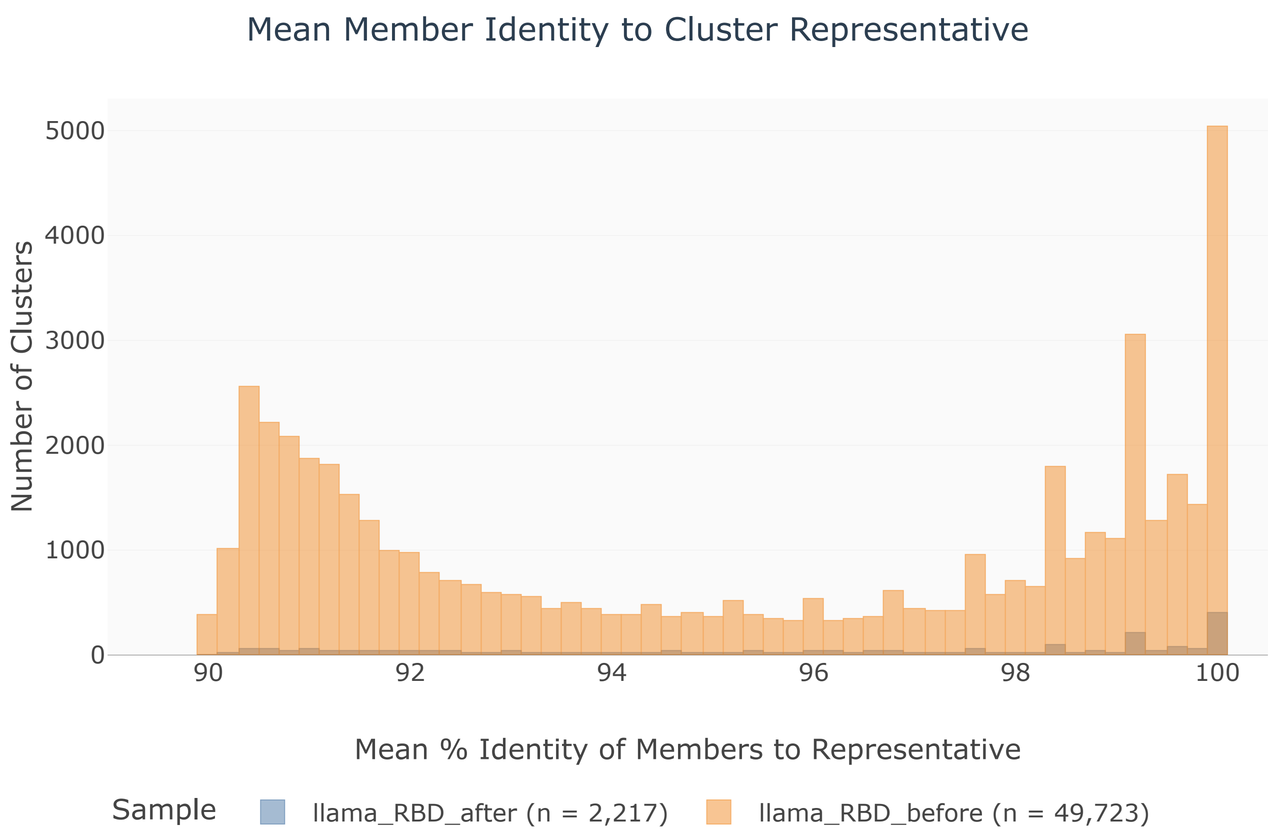

##### Repertoire Diversity Landscape

Bubble chart: Shannon index of the clone abundances (x) against clonality, 1 − Pielou evenness (y); bubble size = unique CDR3 count. Samples move to the lower right as sequences are spread evenly over many clones, and to the upper left as they concentrate into a few dominant lineages. Both axes are computed on cluster sizes, so they are comparable across samples only alongside the depth-normalised richness (clusters per 1000 sequences) reported in the table above.

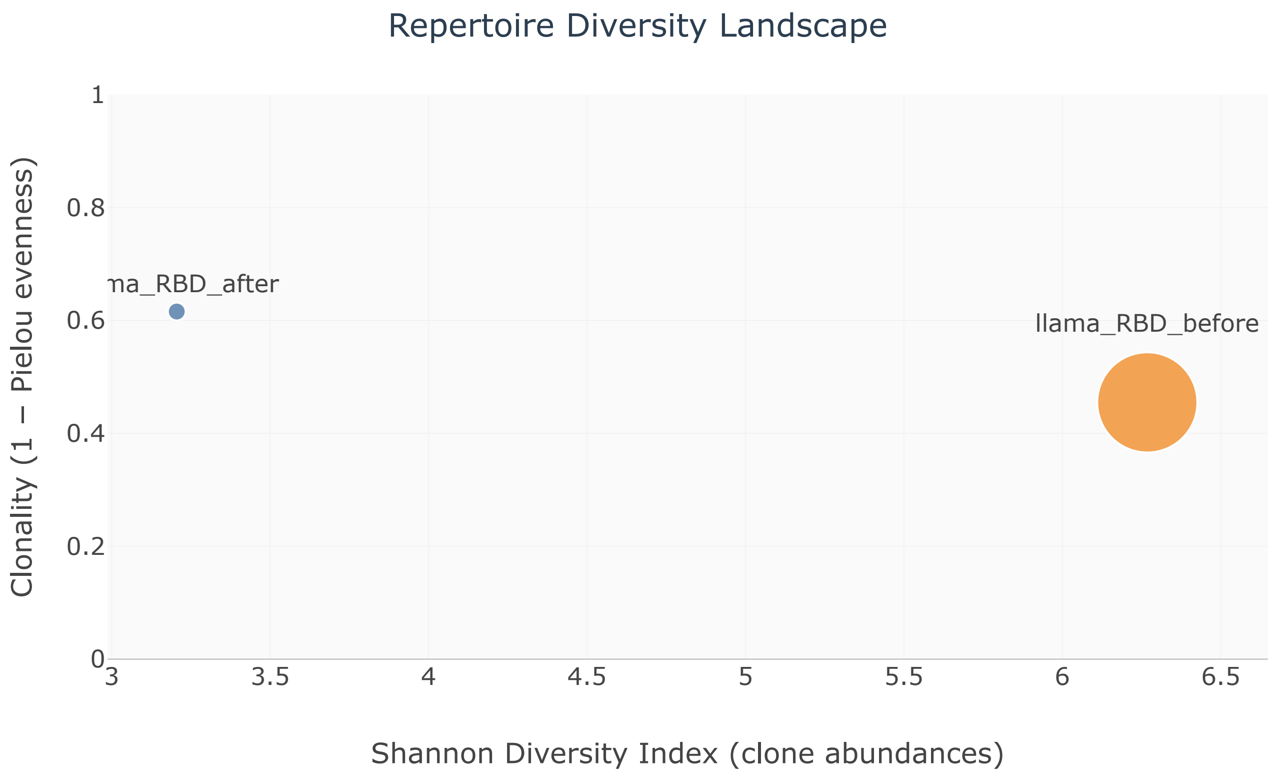


#### 5 Methods Summary

##### 🔬 Pre-processing & Translation

Paired-end FASTQ reads were adapter-trimmed with **Cutadapt** and merged with **FLASH** (max overlap 300 bp). Merged reads were translated in-silico using conserved VHH framework start/end primer motifs (ATG...TVSS pattern; Deschaght 2017).

CDR3 regions were extracted with **nanocdr-x**, a deep learning model trained on camelid VHH sequences.

#### 🧩 CD-HIT Clustering

Translated amino acid sequences were clustered with **CD-HIT**:

- Identity threshold: 0.90 (90%)
- Word size: 5
- Global identity alignment (CD-HIT default `-G 1`); no minimum alignment coverage is imposed
- Four size thresholds analysed: 1, ≥5, ≥100, ≥1000

This follows the validated approach of Deschaght et al. (2017) for large nanobody repertoires.

#### 📐 Diversity Metrics

**Shannon H' (abundance):** $-\Sigma\, p_i \ln p_i$ over the clone frequencies $p_i$ = cluster size / clustered sequences. **Clonality** is 1 − H'/ln(S), with S the number of clusters: 0 when every clone is equally abundant, approaching 1 as the library concentrates into one lineage (undefined, and reported as NA, when S = 1). **Simpson** is the Gini–Simpson index $1 - \Sigma\, p_i^2$, **Gini** the Gini coefficient of the cluster sizes, **D50** the smallest number of clones covering half of the sequences, and **Top1/Top10** the share of the library held by the largest one or ten clones.

**Clusters per 1000 sequences:** richness normalised by sequencing depth. Absolute cluster counts are not comparable between libraries sequenced at different depths; this ratio is.

**CDR3 length evenness (H'):** Shannon entropy of the CDR3 *length* distribution. It measures how evenly paratope lengths are spread and is reported separately from the abundance-based indices above, which it must not be confused with.

**% Expanded Clonotypes:** fraction of clusters with ≥5 members; reflects the proportion of somatically expanded B-cell lineages.

**Intra-clonal homogeneity:** distribution, across clusters, of the mean percent identity of the non-representative members to the cluster representative. Singleton clusters are excluded, the quantity being undefined for them. It is a descriptive summary bounded below by the 90% clustering threshold, not a germline-referenced mutation rate.

#### 📚 Key References


- 📄 Deschaght et al. 2017. *Front. Immunol.* doi:10.3389/fimmu.2017.00420
- 📄 Spinelli et al. 2022. *Front. Immunol.* doi:10.3389/fimmu.2022.927966
- 📄 Muyldermans S. 2013. *Annu. Rev. Biochem.* doi:10.1146/annurev-biochem-071812-105327
- 📄 Bolotin et al. 2015. *Nature Methods*. doi:10.1038/nmeth.3364
- 📄 Desmyter et al. 2002. *J. Biol. Chem.* doi:10.1074/jbc.D200025200
- 📄 Mitchell & Colwell 2018. *Proteins*. doi:10.1002/prot.25497
- 📄 Li & Godzik 2006. *Bioinformatics*. doi:10.1093/bioinformatics/btl158 (CD-HIT)
