## Supplementary Materials for "nanorepertoire: an end-to-end Nextflow pipeline for nanobody repertoire analysis"

*Journal of Immunological Methods*

**Contents**

- S1. Benchmarking dataset details (Table S1)
- S2. Complete overview of the HTML report (Table S2)
- S3. Default parameters and software configuration (Table S3)
- S4. Output data dictionary (Table S4)
- S5. CDR3 annotation quality control (Table S5)
- Supplementary File S1. Complete interactive report (provided separately)

### S1. Benchmarking dataset details

The dataset used to benchmark the nanorepertoire pipeline consists of SARS-CoV-2 immunised llama VHH libraries, totalling 4,771,793 raw paired-end reads. The raw sequencing data were retrieved from the NCBI Sequence Read Archive (SRA) and were originally published by Xu et al. (2021). The two libraries were sequenced to markedly different depth, which is why the pipeline derives depth-normalised statistics alongside the raw counts.

**Table S1.** Summary of the benchmarking dataset.

| Sample ID | SRA accession | Condition | Raw read pairs | Clustered sequences |
| --- | --- | --- | --- | --- |
| llama_RBD_before | SRR13768393 | Pre-enrichment | 4,376,579 | 3,210,691 |
| llama_RBD_after | SRR13768392 | Post-enrichment | 395,214 | 324,341 |

*Data source:* Xu, J., Xu, K., Jung, S., Conte, A., Lieberman, J., Muecksch, F., et al., 2021. Nanobodies from camelid mice and llamas neutralize SARS-CoV-2 variants. Nature 595, 278–282. https://doi.org/10.1038/s41586-021-03676-z.

### S2. Complete overview of the HTML report

The reporting subworkflow (Subworkflow III) separates statistical aggregation from visualisation, so that new panels can be added without modifying the analytical modules. It generates a comprehensive HTML dashboard containing a per-sample metrics table and eleven distinct interactive visualisations. Fig. 2 of the main text highlights four of these; Table S2 describes the full set.

**Table S2.** Visualisations included in the interactive HTML report.

| # | Panel | Description |
| --- | --- | --- |
| 1 | Clonal expansion profile | Stacked bar chart showing the breakdown of clonotypes by size class (<5, 5–99, 100–999, ≥1000 members) per sample. |
| 2 | Cluster abundance at size thresholds | Log-scale grouped bar chart of the cumulative number of clonotypes exceeding each membership threshold (total, ≥5, ≥100, ≥1000). |
| 3 | Top-10 largest clonotypes per sample | Horizontal bar chart identifying the ten most abundant clonotypes per sample, each keyed to the identifier of its representative sequence in clusterbig.csv. |
| 4 | Unique CDR3 sequences per sample | Bar chart of the absolute count of distinct CDR3 paratopes detected per sample. |
| 5 | CDR3 length distribution (violin) | Violin plots with embedded box plots, showing the median, interquartile range and full spread of CDR3 amino-acid lengths per sample. |
| 6 | CDR3 length frequency profile | Filled line chart mapping the frequency of each CDR3 length over the 0–45 residue range, better suited than the violin to resolving multiple modes. |
| 7 | CDR3 boundary offsets | Log-scale bar chart of the residue offsets between the CDR3 called by nanoCDR-X and the motif-based definition, per sample and per terminus. Quality control of the annotation step (see S5). |
| 8 | Amino-acid frequency heat map | Heat map of residue frequency as a percentage of all CDR3 positions, samples as rows and residues as columns. |
| 9 | Side-by-side amino-acid usage | Grouped bar chart allowing direct per-residue comparison of amino-acid usage between samples. |
| 10 | Mean member identity to cluster representative | Histogram of the mean percent identity of the non-representative members of each cluster to its representative, one value per cluster. Singleton clusters are excluded, the quantity being undefined for them. The distribution is bounded below by the clustering threshold, so it describes intra-clonal homogeneity only within that window; it is a descriptive summary and not a germline-referenced somatic hypermutation rate. |
| 11 | Repertoire diversity landscape | Bubble chart plotting the Shannon index computed on clone abundances against clonality (1 − Pielou evenness), with bubble area proportional to the number of unique CDR3 paratopes. |

In addition, the first section of the report presents a per-sample metrics table reporting clonotype richness, clustered sequences, depth-normalised richness (clonotypes per 1,000 sequences), Shannon and Gini–Simpson indices on clone abundances, clonality, the Gini coefficient, D50, the clonal fractions held by the largest and by the ten largest clonotypes, the Shannon entropy of the CDR3 length distribution, and the proportion of expanded (≥5 members) and dominant (≥1000 members) clonotypes. Metrics that are undefined for a given sample, such as evenness and clonality when a sample yields a single cluster, are reported as NA rather than as a substituted value.

### S3. Default parameters and software configuration

The pipeline integrates several tools and standardises their execution within isolated environments — containers (Docker, Singularity/Apptainer) or, as a fallback, Conda — to ensure reproducibility.

**Clustering parameters.** Amino-acid sequence clustering is performed with CD-HIT. The sequence identity threshold (--cdhit_identity) defaults to 90% (-c 0.9) and the word size (--cdhit_word_size) defaults to 5 (-n 5); both are exposed as pipeline parameters and are quoted verbatim in the generated report, so that the document always describes the analysis actually performed. Clustering uses CD-HIT’s default global identity alignment (-G 1), appropriate for full-length VHH sequences; no minimum alignment coverage (-aL / -aS) is imposed.

**Software versions.** The exact versions used for the benchmarking run are controlled by the pipeline configuration.

**Table S3.** Tools and versions used by the pipeline in the benchmarking run.

| Tool | Version | Reference |
| --- | --- | --- |
| Nextflow | 25.04.0 | Di Tommaso et al. (2017) |
| FastQC | 0.12.1 | Andrews (2010), https://www.bioinformatics.babraham.ac.uk/projects/fastqc/ |
| Cutadapt | 4.9 | Martin (2011) |
| FLASH | 1.2.11 | Magoč and Salzberg (2011) |
| CD-HIT | 4.8.1 | Li and Godzik (2006) |
| seqtk | 1.4 | https://github.com/lh3/seqtk |
| Biopython | 1.78 | Cock et al. (2009) |
| nanocdr-x | 1.0.0 | Bagordo et al. (2026) |
| MultiQC | — | Ewels et al. (2016) |
| Quarto | 1.8.27 | https://quarto.org |
| Python | 3.10.20 | — |

**nanoCDR-X execution.** The nanocdr-x module uses a deep-learning model for the precise extraction of CDR3 regions from translated nanobody sequences. To ensure maximum portability without requiring specialised hardware, model inference is optimised to run efficiently on standard CPUs, allowing the pipeline to scale across generic cloud instances and standard HPC environments. For institutions with compatible infrastructure, GPU acceleration can be enabled natively through Nextflow directives.

### S4. Output data dictionary

To facilitate custom downstream analysis, the pipeline outputs structured tabular data and serialised R objects alongside the interactive HTML report.

**Table S4.** Data dictionary of core pipeline outputs.

| File / object | Description and key content |
| --- | --- |
| nanorepertoire_report.html | Interactive HTML dashboard containing the eleven panels of Table S2 and the per-sample metrics table. |
| analysis_report.html | Static scientific summary report rendered with Quarto. |
| nanobodies_report.RData | Serialised R workspace for downstream analysis. |
| clustercounts.csv | Per-sample cluster summary: Clusters, Clusters_of_5, Clusters_of_100, Clusters_of_1000, plus Total_sequences, Top1_count and Top10_count, the denominators required to compute clonal fractions. |
| clusterbig.csv | One row per clonotype: Representative, Count, Identity and Identity_n. Identity is the mean percent identity of the non-representative members to the representative and is NA for singleton clusters, for which it is undefined; Identity_n gives the number of members contributing to that mean. Downstream consumers must exclude NA rather than coerce it to a number. |
| cdrcounts.csv | Total count of unique CDR3 paratopes detected per sample. |
| cdrhists.csv | Frequency distribution (histogram) of CDR3 amino-acid lengths. |
| fastaSeq.csv | Complete table of CDR3 sequences extracted from translated nanobodies, with the full translated sequence and a uniqueness flag. |
| cdr3_boundary_qc.tsv | Per-sample summary of the CDR3 annotation quality control: sequences compared, exact concordance, median and interquartile offsets at each terminus, and the counts of sequences that could not be compared. |
| cdr3_boundary_offsets.tsv | Full distribution of the CDR3 boundary offsets, per sample, terminus and offset value. |
| cdr3_boost_overview_table.tsv | Overview table summarising CDR3 diversity across immunisation and boost metadata. |
| sampledata.tsv | Internal tabular representation of sample metadata (sample ID, individual, immunisation, boost). |
| multiqc_report.html | Aggregated quality control and pre-processing metrics (FastQC, Cutadapt, FLASH). |

### S5. CDR3 annotation quality control

CDR3 boundaries are called by nanoCDR-X. Because every length and composition statistic in the report is conditional on where those boundaries fall, each run re-derives every CDR3 with the motif-based definition used previously for nanobody repertoires (Deschaght et al., 2017) — the loop starting after the framework-3 YYC anchor and ending before the framework-4 WGQ anchor — and compares the two annotations residue by residue. Reads in which the conserved framework-4 tryptophan is substituted are anchored on the C-terminal TVSS motif instead, which places the boundary at the same position.

**Table S5.** Agreement between nanoCDR-X and the motif-based definition.

| Metric | llama_RBD_before | llama_RBD_after |
| --- | --- | --- |
| Sequences with a nanoCDR-X call | 92,744 | 1,687 |
| Comparable (both anchors located) | 86,372 | 1,563 |
| Exact agreement | 99.3% | 99.2% |
| Median offset, N-terminus | 0 | 0 |
| Median offset, C-terminus | 0 | 0 |
| Median called length (residues) | 17 | 18 |
| Median reference length (residues) | 17 | 18 |
| Not comparable, strict WGQ anchor | 15.0% | 12.9% |
| Not comparable, with TVSS fallback | 6.9% | 7.4% |

Where both methods produce a call, the agreement is essentially complete and the median offset is zero at both termini, so the composition and length statistics are not distorted by framework residues leaking into the annotated loop. The two approaches differ in coverage: applied strictly, the motif rule cannot be evaluated on 15.0% and 12.9% of the sequences that nanoCDR-X annotates, because the conserved framework-4 tryptophan is substituted in a substantial minority of reads. Admitting the TVSS fallback recovers most of these, but 6.9% and 7.4% remain inaccessible to a motif definition, almost entirely for want of a locatable framework-3 anchor. Sequences on which the motif rule cannot be applied are counted separately throughout rather than scored as agreements.

### Supplementary File S1. Complete interactive report

Static rendering of nanorepertoire_report.html, the interactive report produced automatically by the pipeline for the benchmarking run described in Table S1. The interactive version is distributed with the pipeline output. This file is provided separately, as a PDF.
